# Temporal origin of mouse claustrum and development of its cortical projections

**DOI:** 10.1101/2022.05.20.492804

**Authors:** Anna Hoerder-Suabedissen, Gabriel Ocana-Santero, Thomas H. Draper, Sophie A. Scott, Jesse G. Kimani, Andrew M. Shelton, Simon J.B. Butt, Zoltán Molnár, Adam M. Packer

**Affiliations:** Department of Physiology, Anatomy and Genetics; Sherrington Building, University of Oxford, Parks Road, OX1 3PT, Oxford, United Kingdom

**Keywords:** claustrum, development, birth date, connectivity, Nurr1/Nr4a2

## Abstract

The claustrum is known for its extensive connectivity with many other forebrain regions, but its elongated shape and deep location have made further study difficult. We have sought to understand when mouse claustrum neurons are born, where they are located in developing brains and when they develop their widespread connections to cortex. We established that a well-characterised parvalbumin-plexus, which identifies the claustrum in adults, is only present from postnatal day (P)21. A myeloarchitectonic outline of the claustrum can be derived from a triangular fibre arrangement from P15. A dense patch of Nurr1+ cells is present at its core, and is already evident at birth. BrdU-birthdating of forebrain progenitors reveals that the majority of claustrum neurons are born during a narrow time window centred on embryonic day (E)12.5, which is later than the adjacent subplate and endopiriform nucleus. Retrograde tracing revealed that claustrum projections to anterior cingulate (ACA) and retrosplenial cortex (RSP) follow distinct developmental trajectories. Claustrum-ACA connectivity matures rapidly, and reaches adult-like innervation density by P10, whereas claustrum-RSP innervation emerges later over a protracted time window. This work establishes the timeline of claustrum development, and provides a framework for understanding how the claustrum is built and develops its unique connectivity.

## Introduction

The claustrum is an elongated, bilateral grey matter structure embedded between the insular cortex and the striatum. Its structure was first described over 200 years ago (Vicq d’Azyr 1786), but its function remains elusive, largely because of its experimentally inaccessible location, and the scarcity of claustrum-specific human brain lesions (Atilgan et al. 2022). The claustrum is widely interconnected with the cortex (Torgerson et al. 2015; Wang et al. 2017), and single claustrum neurons have the largest number of target regions of all neurons studied (Peng et al. 2021). The functional roles proposed for the claustrum all rely on this high degree of connectivity, and range from an involvement in the generation of sleep relevant patterns of activity to top-down executive control. In mice, claustrum neurons are active during slow-wave cortical activity (Narikiyo et al. 2020), and sharp-wave ripples during slow-wave sleep are generated at the anterior medial pole of the dorsoventricular ridge, the reptilian claustrum equivalent (Norimoto et al. 2020). Additionally, REM-sleep active neurons have been reported in the rat and mouse claustrum (Renouard et al. 2015; Maciel et al. 2021). The claustrum has also been reported to function as part of top-down executive control systems involving anterior cingulate cortex (ACA; White et al. 2018), and to contribute to distraction resilience (Atlan et al. 2018). Many of the above behavioural states undergo drastic changes during development, and so far we understand relatively little about emergent brain-wide signalling. The claustrum has extensive connectivity in adults that is ideally placed to orchestrate brain-wide signalling. This makes it an interesting target to study during development. In order to study the developing claustrum we need to be able to reliably find it in young brains, and to differentiate it from nearby structures which may have different functions. The adult claustrum can be delineated by a variety of strategies, including retrograde tracing (Zingg et al. 2018; Marriott et al. 2021), fibre- and cytoarchitecture (Celio 1990; Druga et al. 1993; Dávila et al. 2005; White et al. 2018), and molecular markers (Montiel et al. 2011; WZ Wang et al. 2011, Q Wang et al. 2017), or a combination of these (Grimstvedt et al. n.d.; Wang et al. 2022). Many of the claustrum molecular markers are not detectable in young mouse brains, or appear insufficiently selective to the claustrum when their distribution is assessed from publicly available databases (Bruguier et al. 2020). However some, such as the orphan nuclear receptor 4A2 (*Nr4a2*)/Nurr1, have been used to identify the claustrum in neonatal rat brains (Fang et al. 2021). Retrograde tracing from retrosplenial cortex (RSP) in adult mice is increasingly used as a strategy to identify the claustrum (Zingg et al. 2018; Marriott et al. 2021; Shelton et al. 2022). However, retrograde tracing has never been explored as a labelling strategy in developing brains. It cannot be used before the onset of connections, of course, and thus precludes study of the earliest stages of claustrum development. Moreover, we do not yet know whether claustro-cortical connections form with precision from the start or whether neighbouring regions have transient connectivity to RSP, which would render retrograde tracing a less reliable method in young brains.

Fibre- and cytoarchitecture changes rapidly in young postnatal brains, a period during which neurons finish their migration and axonal connections are first established. In many large-brained species, the claustrum is neatly sandwiched between the external and extreme capsules. While the latter is not easily identified with classical histological stains in the mouse brain, it has been reported that the claustrum is surrounded by densely myelinated fibres (Grimstvedt et al. n.d.; Wang et al. 2017, 2022). Myelinated fibre distribution may thus be a suitable strategy for outlining the claustrum during development, although myelination of the mouse cortex only begins during the second postnatal week (Korrell et al. 2019).

The adult claustrum sends widespread projections preferentially to ipsilateral cortical areas (Wang et al. 2017; Zingg et al. 2018), targeting different laminae in different areas (Wang et al. 2017, 2022). It has strong reciprocal connections with prefrontal and cingulate cortex as well as temporal and retrohippocampal areas (Carman et al. 1964; Kowiański et al. 1998; Vertes 2004; Hoover and Vertes 2007; Atlan et al. 2017; Wang et al. 2017; White et al. 2018; Zingg et al. 2018), suggesting the claustrum may play a crucial role in a variety of cognitive processes. It is thought to be involved in higher functions, and proposed roles include a contribution to inducing cortical down-states during slow-wave sleep, multi-sensory integration during wakefulness or attentional load allocation and salience detection (reviewed in Smith, Lee, & Jackson, 2020). All of these undergo drastic changes during development, and – if the claustrum is indeed a necessary signalling hub - will rely on functional claustro-cortical connectivity. Thus, determining when these connections emerge during development, and linking this time-frame to developmental alterations in the above behaviours, may help to shed light on the function of the claustrum.

During very early development, before the onset of sensory input, cortical networks are synchronised through spontaneous activity in the subplate (Molnár et al. 2020). Previous studies have attempted to establish whether claustrum is the lateral continuation of the cortical subplate, which contains some of the earliest born neurons of the cerebral cortex, revealing a complex picture of both similarities and differences in gene expression between these two structures (Bruguier et al. 2020). In rats, several groups have investigated when claustrum neurons are generated, resulting in a range of likely birth dates from embryonic day (E)13.5 to E15 (Bayer and Altman 1991; Fang et al. 2021), but to the best of our knowledge the equivalent birth dates for mouse claustrum neurons have not been determined. The range of birth dates established for rat claustrum includes the range suggested for subplate, but is equally consistent with a birth-date more similar to cortical layers 5 and 6a. Thus, we aim to establish the birth-date of mouse claustrum neurons with more precision.

Here, we set out to delineate the developing mouse claustrum using molecular and histological means. We identified Nurr1 – a known marker of subplate neurons – as an effective label throughout young postnatal ages but then best combined with myelin staining for clearer delineation of the claustrum in juvenile brains. We further used the myelin-based delineation of claustrum in combination with BrdU labelling to birthdate claustrum, ventral endopiriform nucleus and piriform cortex neurons at P21. Combining a Nurr1 and myelinated fibre-based claustrum delineation strategy with retrograde tracing, we determined the onset of claustrum projections to retrosplenial and anterior cingulate cortex to be during the second postnatal week.

## Methods

### Animals

All animal experiments were approved by a local ethical review committee and conducted in accordance with personal and project licenses under the U.K. Animals (Scientific Procedures) Act (1986). Mice were housed in a temperature-controlled room under a 12 hours light/12 hours dark cycle, with free access to food and water, and pups were kept with their dam until weaning age at P21, or the experimental end-point if earlier.

All brain tissue used was obtained after transcardial perfusion with fixative. Following i.p. injection of an overdose of pentobarbital (200 mg/ml, Pentoject Animalcare), pups or adult mice were perfused, first with 0.1M PBS, followed by 4% formaldehyde (Sigma-Aldrich, F8775) in 0.1M PBS. Brains were dissected out and post-fixed in 4% formaldehyde for a further 24hrs.

### Immunohistochemistry

Free-floating sections were incubated in blocking solution containing 2% donkey serum (Sigma-Aldrich) and 0.2% Triton-×100 (BDH) or 5% donkey serum and 0.5% Triton X-100 (for parvalbumin only) in 0.1M or 0.01M phosphate buffered saline (pH7.4, PBS) for 1-2 hours at room temperature (RT) before being incubated for 48 hours at 4°C with the primary antibodies in the blocking solution. Following the incubation with primary antibodies, sections were washed in PBS before being incubated with secondary antibody in blocking solution for 2 hours at RT. Sections were washed in PBS, counterstained with DAPI (4’,6-diamidino-2-phenylindole, (D3571; Sigma-Aldrich) 5μg/ml in 0.1M PBS) and mounted on microscope slides for imaging. The following primary antibodies were used: goat anti-Nurr1/Nr4a2 (1:100, BioTechne, AF2156), rabbit anti-Cplx3 (1:1000, Synaptic Systems, 122302), rat anti-MBP (1:500, Abcam, ab7349), rabbit anti-parvalbumin (1:500, Swant, P27a), sheep anti-BrdU (1:500, Abcam, ab1893). The following secondary antibodies were used (all at 1:500): donkey anti-goat AlexaFluor594 (Life Technologies, A32758) or donkey anti-goat AlexaFluor568 (Molecular Probes A11057), donkey anti-rabbit AlexaFluor488 (Life Technologies, A21206), donkey anti-rat AlexaFluor488 (Life Technologies, A21208), donkey anti-sheep (Life Technologies, A11016). For anti-BrdU immunohistochemistry (IHC), we used a citrate buffer antigen retrieval protocol, slightly modified from Tang et al., (2007). Antigen retrieval was performed on 3-5 sections/brain by placing them in an Eppendorf tube with 1ml 10mM sodium citrate buffer (pH 6.0), and heating them with a heat-block (Techne) until they reached >97°C (actual temperature of block checked with an alcohol-based glass thermometer). Sections were kept at this temperature for 20 minutes, and subsequently allowed to cool to room temperature. Following washes with 0.1M PBS, the same IHC protocol as above was followed, except that incubation with primary antibody was only 24hrs.

### Birth dating

For birth dating of claustrum neurons, C57/Bl6 or MEC-13-53D mice (Blankvoort et al. 2018, 2020) crossed with homozygous Tg(tetO-GCaMP6s)2Niell (GCaMP6s) mice or Nkx2.1Cre;Z/EG mice crossed with homozygous B6.Cg-Gt(ROSA)26Sortm9(CAG-tdTomato)Hze/J (Ai9) mice were mated and plug-checked daily. Day of plug was designated as embryonic day (E) 0.5. Pregnant dams received a single i.p. injection of 100mg/kg bromodeoxyuridine (BrdU) in sterile saline (BD Biosciences, Oxford, UK) on either E10.5, E11.5, E13.5, or E14.5 (C57 only), E11.5, E12.5 and E13.5 (MEC-13-53D;GCaMP6s only) or E12.5 (Nkx2.1-Cre;Ai9 only). Pups were born between gestational age E19.5 to E21.5 based on plug date, and perfusion fixed (see above) at P21.

Hemispheres from three brains of each litter were sectioned to 50µm coronally using a vibrating microtome (Leica, VT1000S). Sections near the anterior commissure midline crossing were used for immunohistochemical staining (see above). Slices were imaged throughout the section depth in the region of claustrum and dorsal endopiriform/piriform nucleus using a laser scanning confocal microscope in tile-scan and z-stack mode.

For data analysis, an oval region of interest (ROI) was selected to overlap with the MBP-sparse region of the claustrum, the DAPI-dense dorsal region of the piriform cortex, and within dorsal endopiriform nucleus. The same size and shape ROI was used for all regions and all images, and ROIs were selected using only information from the MBP and DAPI channel. Fully-labelled BrdU+ cells within each ROI were manually counted using the FIJI cell counter tool (Schindelin et al. 2012). None of the brains used in this analysis were tdTomato+ (*Nkx2*.*1-Cre;Ai9* animals), and the GCaMP6s signal was not further analysed.

### Connectivity tracing

For connectivity tracing of claustrum neurons using carbocyanine dyes, mouse pups were perfusion fixed at various postnatal ages as described above. Mice of the following strains were used: Rbp4-Cre;Snap25-flox;Ai14 (Cre-negative, or Snap25-fl/+ only, aged P2 and P8), Vglut2-iresCre;Ai9 (Cre+ and Cre-, aged P2, P6 and P10), CTGF-GFP (P2). The diversity of mouse strains was used to determine whether any of them could aid claustrum delineation during development, but the GFP-signal was not detectable after dye incubation, and the *Vglut2-iresCre;Ai9* and *Rbp4-Cre;Ai9* signal was present too broadly, even before dye incubation. For connectivity tracing of claustrum neurons using cholera-toxin B (CTB) injections into live animals, C57 and CD1 mice were used at various postnatal ages.

#### Carbocyanine dye tracing

dye crystals were placed in either ACA, or RSP of each half-brain (hemisphere). DiI (1,1’-dioctadecyl-3,3,3’,3’-tetra-methylindocarbocyanine perchlorate; D3911, Invitrogen) crystals were used for the majority of the brains, and DiA (4-(4-dihexadecylaminostyryl)-N-methylpyridinium iodide; D-3883, Molecular Probes) crystals were used for the few brains expressing tdTomato. A blunt wire tool was used to make a small hole in the surface of the brain at the placement site, based on anatomical surface landmarks, and the crystal pushed into this hole. Brains were incubated in PBS with sodium azide (0.05%) for 4–8-weeks at room temperature or 37°C. Adequate length of incubation for each brain size was confirmed by presence of back-labelled cells in the thalamus.

Following incubation, brains were sectioned to 70μm coronally, using a vibrating microtome. Every 5^th^ section was counterstained with DAPI, before being mounted and imaged. Crystal placement site was confirmed for each brain, and all brains with on-target placements or crystals in immediately adjacent cortical regions were included in this analysis. The region of the claustrum was identified based on nearby anatomical landmarks visible in the DAPI counterstained sections. Back-labelled cells in the claustrum region were counted throughout the section thickness.

#### CTB tracing

51 C57BL/6 postnatal mice from 11 different litters were used for CTB injections into retrosplenial cortex (RSP, age range P8-P79). Similarly, 65 CD1 postnatal mice from 9 different litters were used for CTB injections into anterior cingulate cortex (ACA, age range P1-P34). We used CD1 mice instead of C57 for ACA injections as this resulted in significantly better pup survival following surgery at young ages (26/41 C57 and 44/45 CD1 pups aged <P21 at time of surgery survived for at least 2-day post-surgery). Following surgery, pups were kept with their dam until perfusion or weaning age (P21).

1-2 days prior to surgery, cotton buds soaked in ChloraPrep and covered with Vetbond were placed in the cages, to prepare the dams for the smell. For the CTB-injection surgery in mice >P7, induction of anaesthesia was achieved with 5% isoflurane (IsoFlo, Zoetis) at a rate of 1 L/min in an induction chamber. Then, animals were transferred into a face mask with isoflurane flow which was adjusted to keep them deeply anaesthetised. Mice at or older than P8 were intraperitoneally injected with pre-op Vetergesic (I.P. 0.1 mg/kg - buprenorphine; opioid analgesic) and Metacam (I.P. 5mg/kg, Meloxicam; NSAID), and head-fixed in a stereotaxic frame (Stoelting Digital Stereotaxic). Following skin disinfection with ChloraPrep and skin incision, the bregma-lambda distance was measured and the adult mouse brain Allen Reference Atlas (Allen Institute for Brain Science 2004) coordinates for the injection site were scaled according to the bregma-lambda distance measured for each developing mouse skull (un-scaled coordinates for RSP: AP = -3, ML = 0.5, DV = -1; for ACA: AP = +1.34 ML = +0.3 DV= -1.25 for a mouse with a 4.25 bregma-lambda distance). A hole was drilled at the appropriate coordinates and, after checking correct flow, a glass pipette loaded with CTB-AlexaFluor647 (0.1% wt/vol PBS, Invitrogen Thermo Fisher Scientific, C34778) was inserted. For pups <P8 at injection, surface landmarks were used to target the injection. Injections used sharp pulled glass micropipettes, capable of piercing through the skin and skull. 80 nL of CTB were injected over one minute, and the pipette left in place for at least a further three minutes. After pipette retraction, the skin wound was closed with a small amount of Vetbond for pups, or sutured with coated vicryl for post-weaning mice, before mice were allowed to recover. Pups were returned to the home cage and dam in batches once all animals were fully recovered. Mice with re-opened head-wounds due to maternal overgrooming had the surgery wound re-closed once if the wound was otherwise clean, or were culled and excluded from further analysis if the time-point of culling was <2 days after injection. CTB-injected pups were perfusion fixed either 2 or 5 days after CTB injection, as described above, but using 4% formaldehyde in 0.01M PBS (pH 7.4, Sigma-Aldrich).

Brains were sectioned to 50 µm coronal slices using a vibrating microtome. A series of sections spaced 500 µm apart (for injection ages >P7) or 250 µm (for injection ages <P7), and spanning the entire rostro-caudal extent of the claustrum were used for immunohistochemical labelling (see above) and image acquisition for quantification of CTB-labelled cell counts.

The injection site was confirmed for each brain included in this analysis, by comparing the location of the bulk of the CTB deposit and/or the injection needle track with the adult mouse brain Allen Reference Atlas (Allen Institute for Brain Science 2004). Slices within the region of the injection site were DAPI counter-stained before mounting and imaging. The injection site was confirmed by imaging either with a two-photon-microscope (Bruker) equipped with a Chameleon Vision-S laser (Coherent), imaging at 800 nm for CTB-Al647 and 765 nm for DAPI and using a 16x water immersion objective, with an LSM710 laser scanning confocal microscope, or a Leica epifluorescence microscope equipped with a CY5 filter cube. Images were acquired using Prairie View software (Bruker), ZEN software (Zeiss), or a Leica DFC500 camera with FireCam software. Only brains with clear CTB-Al647 signal along a linear track extending into ACA or RSP, and/or CTB deposits including at least some ACA or RSP were included for further analysis. Brains were excluded from further analysis, if the injection needle track could be located due to tissue damage, but no CTB-Al647 was present at the injection site. Brains were also excluded if CTB-Al647 was found at the injection site, but many CTB-containing blood vessels were present at some distance from the injection site. Images of the claustrum region were obtained with a Zeiss (LSM710) or Olympus (Fluoview F1200) confocal laser scanning microscope. Z-stacks throughout the depth of each slice in the region of claustrum defined by Nurr1 and/or MBP were acquired with an interval of 3.5µm or less. CTB+ cells were manually counted in each of the acquired image stacks, using the FIJI cell counter tool (Schindelin et al. 2012). Only CTB+ cells in the region of the claustrum as defined by the dense Nurr1+ patch (see below) were included in further analysis. Epifluorescence images were aligned with the adult mouse Allen Reference Atlas (Allen Institute for Brain Science 2004) to generate comparable anterior-posterior positions between brains of different sizes and ages, for the purposes of analysing anterior-posterior gradients in retrogradely labelled cell densities. Only cell counts at the level of the striatum were included in further analysis.

### Statistical analysis

All statistical analysis was performed using GraphPad Prism version 9.0.0. Nurr1+ patch size and density were analysed using ANOVA (main effects model) followed by Tukey’s multiple comparison test for anterior-posterior position and brain age, with each brain contributing three or four data points along the AP axis. Individual data points in the graph represent one brain slice each. BrdU+ cell counts were analysed using ANOVA (main effects model) and Tukey’s multiple comparison test for significant differences. Cell counts were averaged for three sections/brain and three brains/litter, such that the individual n was based upon a litter, not sibling pups within a litter. Individual data points in graphs are the mean for each litter. Connectivity tracing timelines were analysed separately depending on method (carbocyanine vs CTB) and location of injection using ANOVA to detect differences between survival times, and Spearman’s rank correlation to determine correlations between injection age and number of cells. Anterior-posterior gradients were analysed separately by injection site using ANOVA and Tukey’s multiple comparison test. Individual brains (i.e. injections) were considered independent, even if animals derived from the same litter. All summary data is reported as mean±sem.

## Results

### Identification of the claustrum in young postnatal mouse brains

Parvalbumin (PV) immunohistochemistry has been widely used to delineate the claustrum because of the dense PV+ plexus of fibres present in the dorsal claustrum complex (Celio 1990; Druga et al. 1993; Wang et al. 2017; White et al. 2018), and is the preferred claustrum marker when comparing different species. PV-immunoreactivity, including presence of the dense fibre plexus has previously been reported to reach adult-like distribution and density in the P23 mouse claustrum, but only faintly stained cell-bodies with no clear boundary to surrounding structures are evident at P4 (Dávila et al. 2005). We investigated the development of the PV+ fibre plexus in the claustrum at intervening time-points in C57/Bl6 mice (Fig 1). The PV+ plexus is well-established at P21 (Fig 1D; n=3 brains), with clear boundaries to surrounding tissue, although the plexus has not yet reached adult density (c.f. Fig 1C, n=3 brains). At P14, however, very few PV+ fibres were evident in the claustrum region, despite the presence of brightly labelled PV+ cell bodies (Fig 1E; n=3 brains). At this age, PV immunohistochemistry alone is insufficient to delineate the claustrum.

**Figure 1.**
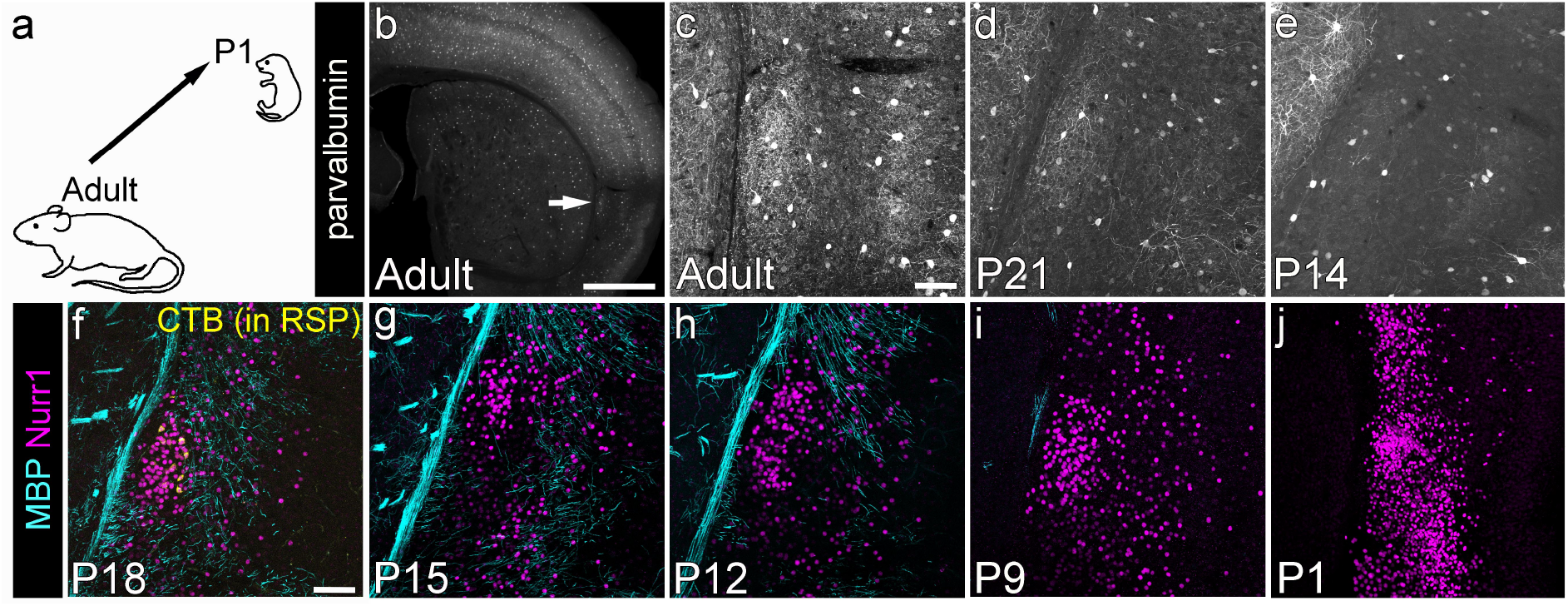
Claustrum labelling in developing (postnatal) brains. (a) we used well-established adult claustrum labelling strategies and determined whether the same labelling can be used to identify the claustrum in developing brains. (b) Epifluorescence image of an adult mouse brain section stained with parvalbumin (PV) to reveal the ‘PV plexus’ commonly used as a claustrum landmark. (c-e) Maximum intensity projection laser scanning confocal microscope images of the claustrum region stained for PV in adult (c), P21 (d) and P14 (e) mouse brains. A loose PV+ plexus is present at P21, but is not detectable at P14, despite the presence of strongly labelled cell bodies. (f-j) Maximum intensity projection of confocal images of the claustrum at different ages, stained for Nurr1 and myelin basic protein (MBP). (f) Claustrum cells can be labelled with retrograde tracer injections into retrosplenial cortex (RSP, as demonstrated here by cholera toxin B-AlexaFluor647 (CTB) labelling (yellow) of cells in claustrum). The region of CTB+ cells is surrounded by dense myelinated fibres in a triangular arrangement, but the claustrum itself contains little myelin. A dense patch of Nurr1+ cells is located within the region of sparse MBP labelling, and overlaps the CTB+ cell cluster at P18. (g-j) The Nurr1+ patch of cells remains visible at all postnatal ages, whereas the MBP-labelling is less pronounced in younger brains and no longer provides an outline of the claustrum at ages younger than P12 (h-j). Note, that in the small P1 brains, the size of Nurr1+ nuclei is also decreased (j). Scale bars = 1mm (b) or 100µm (c-j)

It has been previously reported, that the cell-dense adult claustrum is located “in an oval region that is less well myelinated than its surrounding structures” (Wang et al. 2017). Specifically, the MBP+ fibres meet in a triangular arrangement in or surrounding the claustrum in adult brains (Grimstvedt et al. n.d.). Immunohistochemistry against myelin basic protein (MBP), a component of the mature myelin sheath, confirms that the myelinated fibres are sparse in the claustrum, as identified by retrograde tracing from retrosplenial cortex (Fig 1F). The first MBP-containing myelin in the murine cortex can be detected at the striato-cortical junction at P7 (Korrell et al. 2019), but does not yet extend to the lateral location of the claustrum (data not shown). MBP in the regions surrounding the claustrum is first detectable in the external capsule at P9 in C57/Bl6 mice (Fig 1I, n=3 brains). At this age, only few axons are myelinated, with most of the external capsule remaining unmyelinated. By P12, the external capsule contains many more myelinated axons, and myelinated fibres are starting to surround the claustrum on two sides in an inverted V-shape (Fig 1H, n=5 brains). From P18 (Fig 1F; n=3 brains) the MBP+ fibres surrounding the claustrum begin to resemble a bird’s nest-like arrangement, with only a small region, containing the cluster of RSP-projecting claustrum cells, remaining almost completely unmyelinated. We have used the pattern of MBP immunoreactivity to assist in the localisation and delineation of the claustrum throughout this manuscript, at ages where it is present.

Nurr1/Nr4a2, an orphan nuclear receptor, was initially reported to broadly label lateral cortex, including claustrum, in the adult rat (Arimatsu et al. 2003), and has since been used to label the claustrum complex in neonatal rats (Fang et al. 2021) and adult mice and rats (Montiel et al. 2011; Wang et al. 2011; Niu et al. 2022). Nurr1 immunoreactive nuclei are not restricted to the claustrum at any postnatal age in the mouse brain (Fig 1F-J), but they do form a particularly dense cluster immediately adjacent to the external capsule in the region of the claustrum. The Nurr1+ nuclei are more loosely spaced, less brightly labelled and more distant from the external capsule both dorsal and ventral to the claustrum. At the level of the anterior commissure, this ‘Nurr1+ patch’ of cells falls within the MBP-sparse triangle and overlaps with the cluster of RSP-projecting claustrum cells (Fig 1F), and is probably the equivalent of the ‘ventral claustrum’ in adult mice (Grimstvedt et al. n.d.). Moreover, it is evident already at P1 in the mouse brain (Fig 1J, n=3 brains). The dense expression of Nurr1 can therefore be used from birth to distinguish the claustrum from adjacent structures. Other markers, useful for locating the adult claustrum, only start to provide a suitable label from the end of the second postnatal week at the earliest.

We further examined the anterior-posterior extent of the Nurr1+ patch. The dense Nurr1+ patch of cells in the claustrum is identifiable throughout the anterior-posterior extent of the claustrum, but it varies considerably in size along the AP axis (Fig 2). Anatomical landmarks were used to identify the anterior-posterior level, because distance from bregma varies across development as the brain expands. The Nurr1+ patch first emerges anteriorly as a small cluster of cells where the anterior forceps of the corpus callosum begins to form a horse-shoe shape (Fig 2B,F; n=3 brains each at P9 and P21 respectively). The Nurr1+ patch then reaches its maximum extent at the point when the corpus callosum first crosses the midline (Fig 2C,G; n=3 brains at P9 and P21 respectively). At the level of the anterior commissure midline crossing, the Nurr1+ patch of cells sits within the inverted V-shape of MBP-fibres (Figs 1&2), and/or fits into the centre of the MBP+ ‘bird’s nest’ in more mature brains (n=22 brains across a range of ages from P18-P79, see Fig 1F and Suppl Fig 1). Here the Nurr1+ patch covers a smaller area than at the level of the anterior corpus callosum midline crossing (Fig 2D,H; n=3 brains at P9 and P21 respectively). Further caudally, the Nurr1+ patch of cells continues to decrease in size, and eventually becomes indistinguishable as a distinct patch at the level of the anterior hippocampus when both the upper and lower blade of the dentate gyrus are present. Excluding the most anterior sections where presence or absence and therefore size of the Nurr1+ patch can be ambiguous, we determined the area of the Nurr1+ patch and the cell density of Nurr1 at various ages from maximum intensity projection confocal images (n=22 brains, aged P5-P36, Fig 2N). Each brain is represented at least at three different locations in this analysis, but most are represented at four locations. In these images the Nurr1+ patch is at its biggest anteriorly, and decreases significantly along the AP axis (main effects model ANOVA F (3, 63) = 26.85, p<0.0001; Tukey’s multiple comparison test is significantly different for all comparisons except between very anterior and anterior locations; Fig 2N). Nurr1+ patch size is also affected by brain age (main effects model ANOVA F (9, 63) = 2.764, p=0.0086). This does not cleanly correlate with a Nurr1+ cell density gradient along the AP axis. Although there is a significant effect of anterior-posterior position on Nurr1+ cell density within the claustrum (main effects model ANOVA F (3, 63) = 3.701, p=0.0161; Suppl Fig 2), Tukey’s multiple comparison test indicates that only the Nurr1+ patch density at the approximate level of the anterior commissure midline-crossing is significantly different from the anteriorly adjacent claustrum (Tukey’s multiple comparison test, p=0.0318). There is a strong effect of age on Nurr1+ cell density in the claustrum (main effects ANOVA F (9, 63) = 16.55, p<0.0001), which is exclusively due to the much higher cell density throughout the claustrum in the youngest (and therefore smallest) two brains (Tukey’s multiple comparison test p<0.01 for all comparisons involving P5 and P8 brains). Interestingly, the Nurr1+ patch cross-sectional area is not significantly different when comparing P5 and P8 brains with any other age (Tukey’s multiple comparison test, p=0.57-0.99).

**Figure 2.**
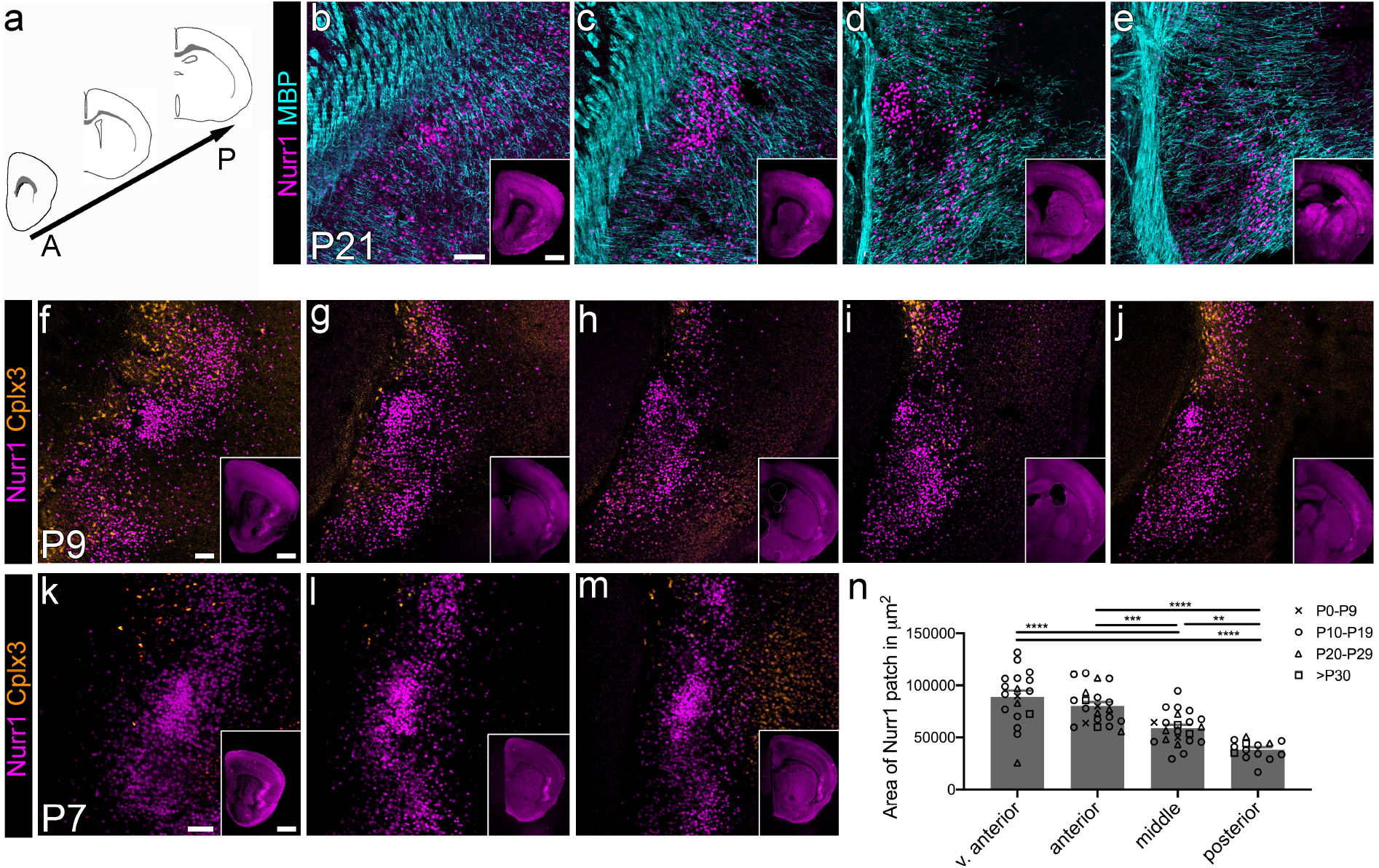
Markers delineating the developing claustrum along the anterior-posterior (AP) axis. (a) We assessed different developmentally suitable claustrum markers along the AP axis. (b-e) Maximum intensity projection confocal laser scanning microscope images of Nurr1 and myelin basic protein (MBP) immunofluorescence at P21 at selected AP levels of the claustrum. AP position indicated by insets (epifluorescence images of Nurr1 signal). Note that the claustrum is always identifiable by the dense patch of Nurr1+ cells, although the size of the patch varies considerably along the AP axis. The diagonal orientation of the MBP+ fibres can help to localise the claustrum at all AP levels, but the MBP-sparse claustrum region is most evident at the level of the anterior commissure (d). (f-j) Maximum intensity projection confocal images of Nurr1 and Cplx3 immunofluorescence at P9 at selected AP levels of the claustrum. AP position indicated by insets (epifluorescence images of Nurr1 signal). The Nurr1+ patch is bordered dorsally by Cplx3+ subplate cells, and ventrally by Cplx3+ cells in the endopiriform nucleus. At very anterior levels some Cplx3+ cells are also present medial to the claustrum (f). (k-m) Epifluorescence images of Nurr1 and Cplx3 immunofluorescence at P7 at selected AP levels of the claustrum (AP position indicated by insets). While the Nurr1+ patch is very distinct at P7, there are hardly any Cplx3+ at the dorsal border of claustrum, and none in endopiriform nucleus. (n) Quantification of Nurr1+ patch cross-sectional area as measured on maximum intensity projection confocal images across a range of postnatal ages. At the most anterior levels, the Nurr1+ patch is very variable in size. There is a significant decrease in Nurr1+ patch size along the anterior-posterior axis (main effects model ANOVA F (3, 63) = 26.85, p<0.0001). Scale bars = 100µm or 1mm (insets). ** p<0.01, *** p<0.001, **** p<0.0001

*Cplx3*, encoding a presynaptic protein, has been reported to label both layer 6b dorsal to the claustrum, and the endopiriform nucleus ventral to the claustrum in adult brains (Grimstvedt et al. n.d.), and may thus help to delineate the claustrum. We observed Complexin 3 (Cplx3)+ immunohistochemical labelling of the subplate at P7, in agreement with previous studies (Hoerder-Suabedissen et al. 2009; Hoerder-Suabedissen and Molnár 2013), but at this age hardly any Cplx3+ cells were found in the endopiriform nucleus (n=3 brains; Fig 2K-M). By P9, Cplx3+ cells demarcate the dorsal edge of the Nurr1+ patch, and therefore the dorsal edge of the claustrum, but are absent from claustrum itself (Fig 2F-J; n=3 brains). At anterior levels, many Cplx3+ cells are found medial to the Nurr1+ patch of claustrum cells, indicating the presence of subplate medial to the claustrum. Due to the limitations in Cplx3 immunohistochemistry at the youngest ages, we used MBP and/or Nurr1 immunohistochemistry for the remainder of this manuscript to delineate the claustrum.

### BrdU labelling of claustrum complex neurons reveals distinct birth dates for claustrum and endopiriform nucleus

We sought to understand when claustrum neurons are generated, to determine how they fit into the general pattern of brain development and neurogenesis of the cortex. We used BrdU birth-dating to determine the peak of neurogenesis in the claustrum in mice (Fig 3). All analysis was performed on sections at the approximate level of the anterior commissure midline crossing in C57/Bl6 mice aged P21, to enable reliable identification of the claustrum based on the myelo-architecture of the surrounding tissue. In common with six-layered neocortex, few neurons in the claustrum complex and piriform cortex are generated at E10.5, although BrdU+ cells were found in other parts of the brain (Fig 3A; six brains from n=2 litters). The peak of neurogenesis for endopiriform nucleus is at E11.5 (Fig 3B; nine brains from n=3 litters), and very few neurons in this region are generated on or after E13.5 (Fig 3D,E; nine brains from n=3 litters). Neurons in the claustrum (as defined here by the MBP-sparse region) are primarily generated at E12.5 (Fig 3C,C’; nine brains from n=3 litters). E12.5 BrdU-label is also dense directly ventral to the claustrum as defined above (not quantified), but no robust histological feature could be identified to demarcate the boundary between the E12.5-generated claustrum, and the E11.5-generated endopiriform nucleus. The peak of neurogenesis for piriform cortex is also at E12.5, but extends for a longer period, with considerable numbers of neurons continuing to be generated at E13.5 (nine brains from n=3 litters) and E14.5 (six brains from n=2 litters). Two-way ANOVA was used to analyze the effect of injection age and brain region on BrdU+ cell density. There was a statistically significant interaction between the injection age and brain region (F (8, 24) = 9.829, p<0.0001) on BrdU+ cell density. Main effects analysis showed that both injection age and brain region had a statistically significant effect on BrdU+ cell density (p < 0.0001 and p=0.0003 respectively). Significantly more neurons in endopiriform nucleus are generated at E11.5 than at any other age (Tukey’s multiple comparison test p<0.005 for all comparisons involving E11.5), with all other ages being non-significantly different from each other. Similarly, significantly more neurons in claustrum are generated at E12.5 than at any other age (Tukey’s multiple comparison test p<0.0001 for all comparisons involving E12.5), with all other ages being non-significantly different from each other. Conversely, the density of BrdU+ neurons in piriform cortex is only significantly different between the earliest injection ages and later time-points, confirming the extended period of piriform neurogenesis, compared to the much shorter time-windows of endopiriform nucleus and claustrum neurogenesis.

**Figure 3.**
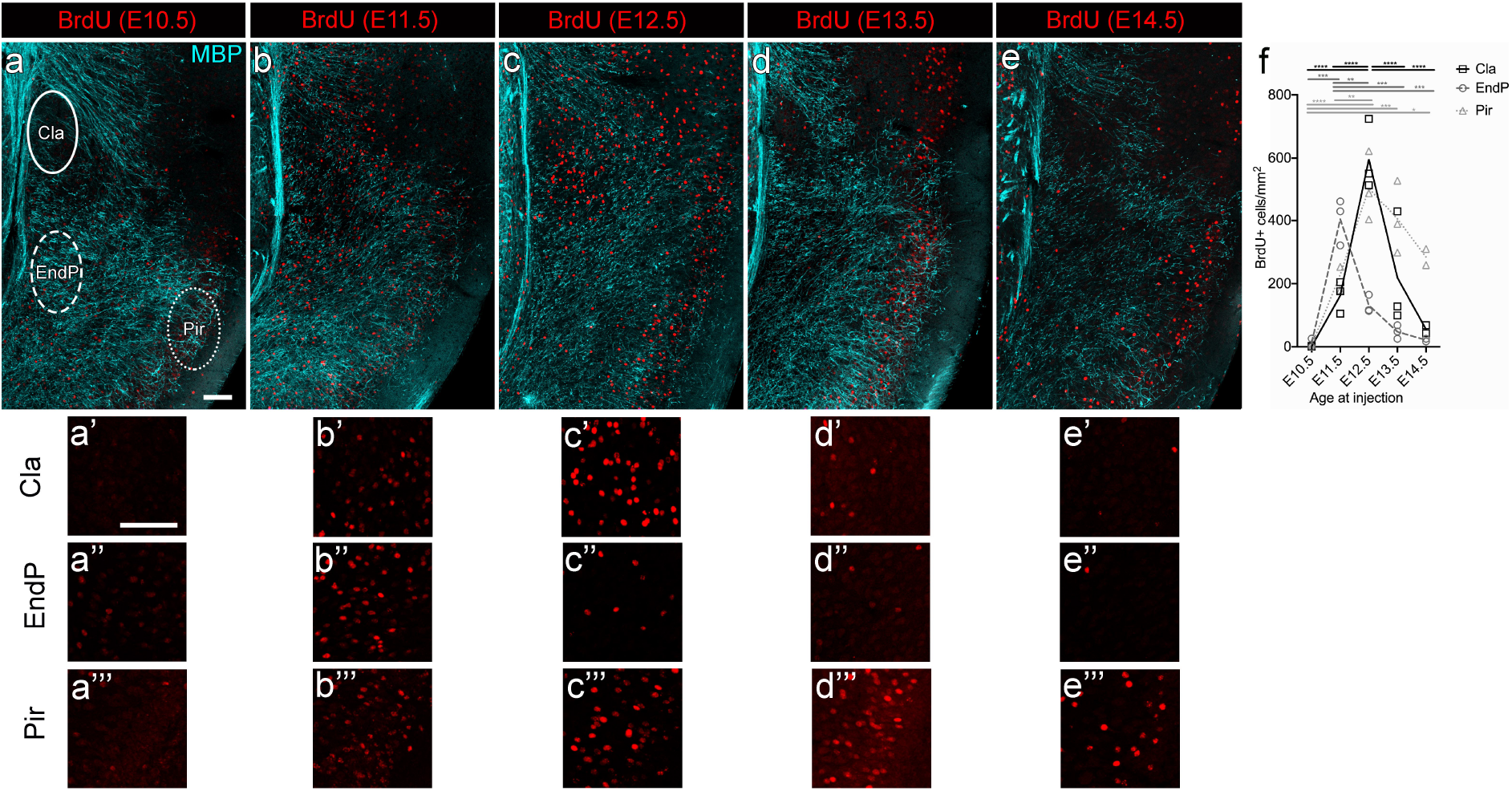
Claustrum neurons have distinct birth-dates from neurons in nearby endopiriform nucleus and piriform cortex. Timed-pregnant mice were injected i.p. with 100mg/kg BrdU at one of E10.5, E11.5, E12.5, E13.5 or E14.5, and brains of pups were collected at P21 before being processed for BrdU-immunohistochemistry. (a-e) Maximum intensity projection confocal laser scanning microscope images of the claustrum region at P21, immunohistochemically stained for BrdU and MBP. All images are taken at the level of the anterior commissure. BrdU+, fully-labelled nuclei were quantified in the claustrum (Cla), endopiriform nucleus (EndP) and piriform cortex (Pir) in the regions outlined in (a). Higher magnification images of the BrdU signal in claustrum (a’-e’), endopiriform (a’’-e’’) and piriform cortex (a’’’-e’’’) are shown underneath. There are few BrdU+ cells present in the claustrum and surrounding brain areas following an E10.5 injection (a, a’). BrdU-injections at E11.5 through to E13.5 result in abundant labelling of cells in lateral cortex, with regional variability in the distribution. Endopiriform nucleus is most heavily labelled following E11.5 injections (b, b’’), whereas claustrum is most densely labelled following E12.5 injections (c, c’). Piriform cortex contains cells born across a range of time-points. (f) Quantification of BrdU+ cell density in the claustrum, endopiriform nucleus and piriform cortex. Each data point is the mean of three brains from the same litter of pups. Scale bars = 100µm. * p<0.05, ** p<0.01, *** p<0.001, **** p<0.000

### Is there a subplate medial to the dorsal claustrum, or is the subplate continuous with the claustrum?

The subplate is typically defined as ‘a thin band of cells adjacent to the white matter and containing some of the earliest born neurons in the cerebral cortex’ in rodents (Angevine Jr. and Sidman 1961; Price et al. 1997; Hoerder-Suabedissen and Molnár 2013). In more mature brains the region containing the remaining early-born neurons, and expressing the same molecular markers, is referred to as layer 6b. Anatomically, its ventral neighbour is the dorsal claustrum. But are they distinct structures or do they share some continuity or contain similar cells?

The subplate and the claustrum complex share some gene expression (Montiel et al. 2011; Wang et al. 2011; Bruguier et al. 2020), and Nurr1 used here as a marker for the earliest claustrum, is expressed in subplate in dorsal cortex, but more broadly in lateral cortex (Arimatsu et al. 2003; Hoerder-Suabedissen et al. 2009). Cplx3+ cells, on the other hand, are restricted to the subplate in both dorsal and lateral cortex (Hoerder-Suabedissen et al. 2009). It is additionally expressed in the endopiriform nucleus, but not in the claustrum (Fig 2 and Grimstvedt et al. n.d.). However, especially in anterior sections, a few scattered Cplx3+ cells can be found embedded in the external capsule, medial to the claustrum (Fig 2). *Connective tissue growth factor* (*Ctgf)*, another commonly used subplate marker, as well as the transgenic strain CTGF-2A-dgCre, show expression similar to Cplx3, i.e. present in subplate/layer 6b and endopiriform nucleus, but absent from most of the claustrum (see the adult Allen mouse brain *in situ* hybridisation database (Lein et al. 2007) and (Grimstvedt et al. n.d.; Montiel et al. 2011; WZ Wang et al. 2011, Q Wang 2017; Erwin et al. 2021)). Brains injected with BrdU at E11.5, showed a clear band of BrdU+ cells in the cortical subplate just above the white matter (data not shown), consistent with subplate being ‘early born’. All E11.5 injected brains (but not every image for each brain) also showed BrdU+ cells within the external capsule medial to the claustrum. The BrdU+ cells were sometimes arranged as a thin band, and always located outside of the area used for claustrum birth-date quantification (see Fig 3B). Notably, the peak of neurogenesis for the claustrum core, the region devoid of Cplx3 or *Ctgf* expression, is one day later (at E12.5). This suggests that there is a subplate medial to the claustrum, rather than the dorsal claustrum being a direct continuation of the subplate.

### Onset of claustral innervation of retrosplenial cortex

We used carbocyanine dye tracing in fixed brains (P2-P10) and cholera toxin B (CTB)-labelling *in vivo* (P8-P74) to determine when claustrum neurons first extend an axon into retrosplenial cortex (Fig 4), one of their adult projection target regions. We only analysed ipsilateral projections, as the RSP-projecting claustrum neurons in mice have been reported to project almost exclusively ipsilaterally (Zingg et al. 2018).

**Figure 4:**
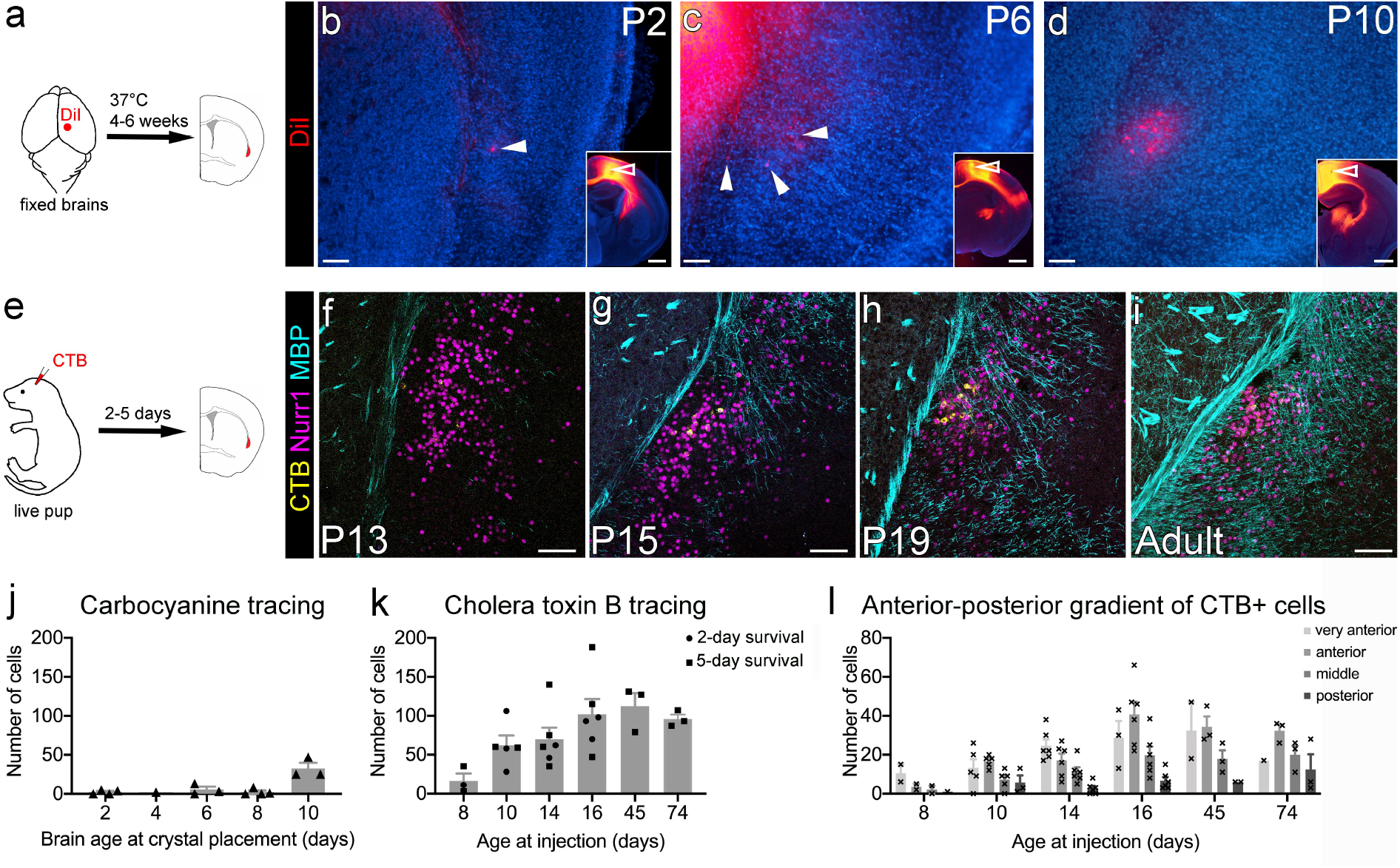
Claustrocortical axons reach retrosplenial (RSP) cortex during the second postnatal week. (a) Carbocyanine dye tracing was used in fixed brains to determine the approximate age at which cells in the claustrum and lateral cortex extend axons to RSP. (c-e) Epifluorescence images of the claustrum region as delineated by relative cell density of DAPI-stained nuclei (blue), and carbocyanine labelling. The crystal placement site for each brain is shown in the insets, with white open arrowheads pointing at the placement site. A few DiI+ cells were found in the region of the claustrum, even at the youngest ages studied (arrowheads in b, c). By P10, a cluster of retrogradely labelled cells was visible in claustrum (d). (e) To determine when adult-like innervation density from claustrum to RSP is reached, we used retrograde labelling with CTB-AlexaFluor647 injected into RSP of live pups and mice, with post-injection survival of two or five days. (f-i) Maximum intensity projection confocal images of CTB-labelled cells (yellow) in the claustrum region as defined by Nurr1 (magenta) and MBP (cyan) immunofluorescence signal. All images are from 5-day post-injection survival, and age of brain at fixation is given for each image. Very few CTB+ cells were found in P8-injected brains (f). (j) Mean±sem of all claustrum cells retrogradely labelled with carbocyanine, counted in every 5^th^ section along the AP extent of the claustrum of a given brain. There was a sharp increase in the number of retrogradely labelled claustrum cells at P10, compared to earlier ages. (k) Mean±sem of all CTB+ retrogradely labelled claustrum cells counted in every 10^th^ section along the AP extent of the claustrum of a given brain. Individual brains shown as triangles (2-day survival) and squares (5-day survival). There was no significant difference in the number of retrogradely labelled claustrum cells between 2-day and 5-day post-injection survival time (ANOVA, F (1, 17)=1.113, p=0.3063), but a strong positive correlation between number of back-labelled claustrum cells and increasing brain age (Spearman’s r=0.9429, p=0.0167; k). (l) Anterior claustrum projects more strongly to RSP throughout development. Retrogradely labelled cell counts in individual slices were binned by anterior-posterior location. ‘Very anterior’ corresponds to slices at and just posterior to the beginning of striatum. ‘Anterior’ corresponds to slices at and just posterior to the first midline crossing of the corpus callosum. ‘Middle’ is at the level of the anterior commissure midline crossing, and ‘posterior’ is caudal to that. More RSP-projecting neurons are located in the anterior claustrum than in the more posterior regions nearer the injection site. This gradient is more pronounced in older brains. There was a significant effect of both age at injection (ANOVA, F (5, 81)=9.618, p<0.0001) and anterior-posterior position (F (3, 81)=22.87, p<0.0001). Scale bars = 100µm and 1mm (insets)

For each brain, a carbocyanine crystal was placed in the approximate location of anterior RSP from the dorsal surface of the intact hemisphere, and hemispheres or whole brains were incubated for 4-6 weeks to allow the dye to diffuse. Location of the injection site was confirmed to include RSP for all brains included in this analysis, but the crystal placement site typically extended into the more lateral secondary and primary motor cortices (MOs and MOp, respectively) as well. For carbocyanine dye placements in RSP/MOs/MOp, there were some brains at each age up to P8 in which we could find no retrogradely labelled cells in the claustrum, despite the presence of back-labelled thalamic cells in the same brains. We also observed carbocyanine labelled neurons in the lateral subplate in all brains, and in older brains also in other cortical layers. In contrast, some of the youngest aged brains used (P2) contained a small number of back-labelled cells in the claustrum, typically in more anterior sections (see Fig 4B; 2/4 brains). At P10, all brains investigated (n=3) contained back-labelled cells in the claustrum, sometimes forming a distinct cluster of cells (see Fig 4D; 3/3 brains, but not every section for each brain). There was a clear increase in the number of RSP-projecting, putative claustrum cells in P8 and P10 brains (Fig 4D,J), but no statistically significant correlation between increasing brain age, and number of back-labelled putative claustrum cells across the entire age spectrum (Spearman’s r=0.67, p=0.2667).

To extend our analysis and focus on the onset of the adult like pattern of innervation, we switched to CTB-AlexaFluor647 injections, a largely retrograde tracer (Conte et al. 2009). CTB was injected into posterior RSP (n=51 mice from 11 litters) based on scaled stereotaxic coordinates, and the injection site was confirmed to include RSP for each brain included in the analysis here (n=26 from 6 litters). CTB-labelling requires injection *in vivo*, and a long-enough post-injection survival time for adequate transport of CTB back to the cell body. To control for possible variability in the duration over which CTB is present in the extracellular space, we used two different post-injection survival times: five days, which we knew to give adequate back-labelling of claustrum cells in adult brains in our hands (Grimstvedt et al. n.d.; Shelton et al. 2022), and two days, to reduce the risk of newly arriving axons being able to take up CTB days after the injection took place. There was no significant difference in the number of retrogradely labelled claustrum cells between 2-day and 5-day post-injection survival time (ANOVA, F (1, 17)=1.113, p=0.3063; Fig 4K). Different post-injection survival times were therefore grouped together for all subsequent analysis.

We observed retrogradely labelled claustrum neurons from our earliest time point, P8. There was a strong positive correlation between the number of back-labelled claustrum cells and increasing brain age (Spearman’s r=0.9429, p=0.0167). Adult-like innervation density was reached around P16.

There was a pronounced anterior-posterior gradient, with many more back-labelled cells found in the anterior claustrum at most ages (Fig 4L). To analyse this further, we binned cell counts into four anterior-posterior positions. There was a significant effect of both brain age at injection (ANOVA, F (5, 81)=9.618, p<0.0001) and anterior-posterior position (F (3, 81)=22.87, p<0.0001). At most ages, the highest number of cells were found in the ‘very anterior’ claustrum, which corresponds to sections at the level of the anterior end of the striatum or the ‘anterior’ claustrum, which corresponds to sections at approximately the anterior-most corpus callosum midline crossing. There was no statistically significant difference between cell counts in ‘very anterior’ and ‘anterior’ claustrum, but both were significantly different from ‘middle’ and ‘posterior’ claustrum (Tukey’s multiple comparison test: very anterior vs anterior p=0.8654, very anterior vs middle p=0.0051, very anterior vs posterior p<0.0001, anterior vs middle p=0.0001, anterior vs posterior p<0.0001, middle vs posterior p=0.0112).

### Establishment of claustro-cortical projections to anterior cingulate cortex

Claustrum is reported to project to ACA with higher density than to RSP in adult mice (Wang et al. 2017; Zingg et al. 2018), but in contrast to RSP, ACA cells also receive inputs from insula and other claustrum-adjacent structures. We used carbocyanine dye tracing in fixed (P2-P10) and CTB-labelling in living brains (P3-P34) to determine when claustrum neurons first extend an axon into anterior cingulate cortex (ACA), and when adult-like connectivity is achieved (Fig 5).

**Figure 5:**
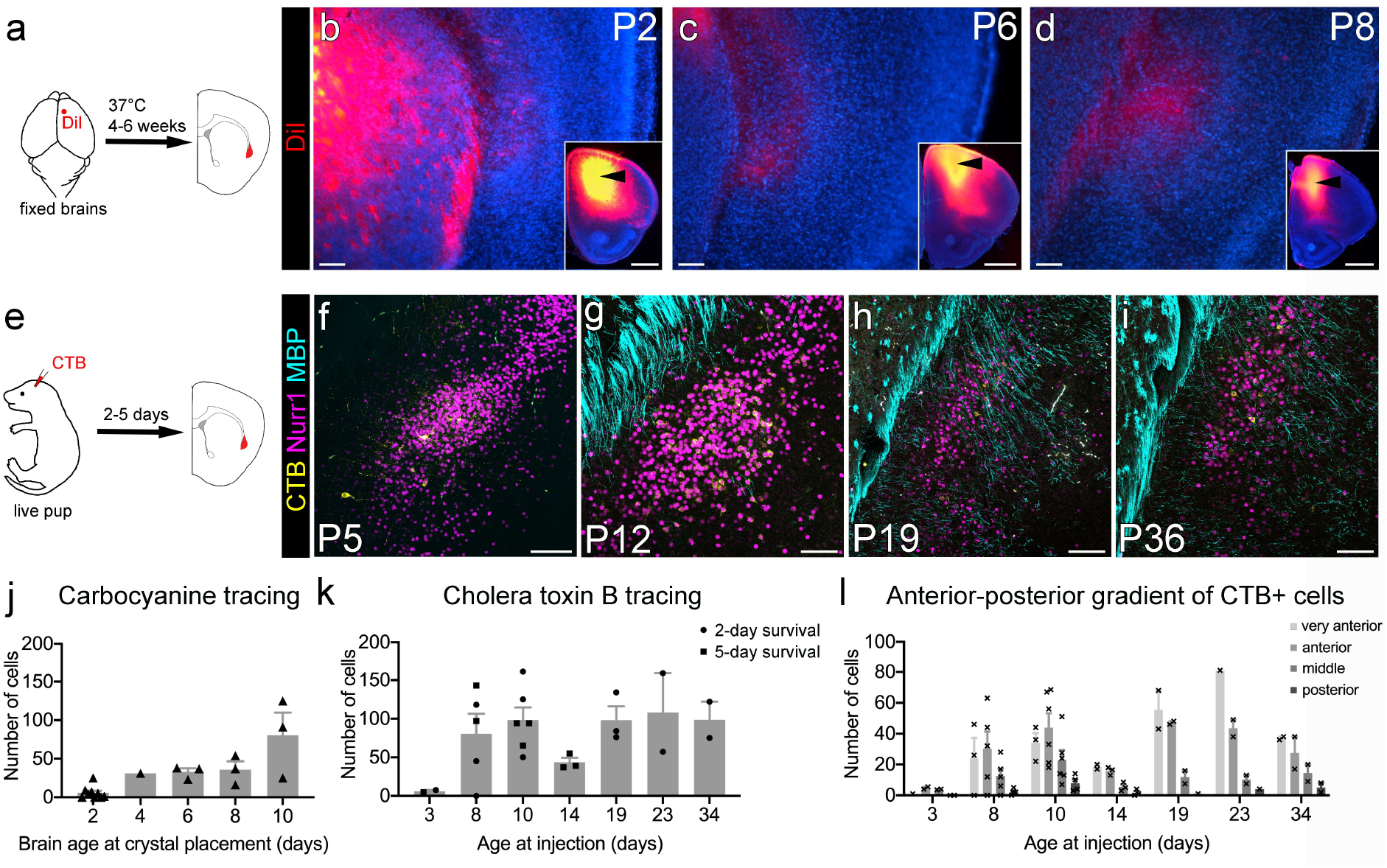
Claustrocortical afferents to anterior cingulate cortex reach peak density during the second postnatal week. (a) Carbocyanine dye tracing was used in fixed brains to determine the approximate age at which cells in the claustrum and lateral cortex extend axons to anterior cingulate cortex (ACA). (b-d) Epifluorescence images of the claustrum region and adjacent lateral cortex as delineated by relative cell density of DAPI-stained nuclei (blue), and carbocyanine labelling. A few cells were found dorsal to the claustrum, and occasionally in the region of the claustrum, even at the youngest ages studied. The crystal placement site for each brain is shown in the insets, with black arrowheads pointing at the placement site. (e) To determine when adult-like connectivity from claustrum to ACA is reached, we used retrograde labelling with CTB-AlexaFluor647 injected into ACA of live pups and mice, with post-injection survival of two or five days. (f-i) Maximum intensity projection laser scanning confocal images of CTB-labelled cells (yellow) in the claustrum region as defined by Nurr1 (magenta) and MBP (cyan) immunofluorescence signal. All images are from 2-day post-injection survival, and age of brain at fixation is given for each image. Very few CTB+ cells were found in the claustrum in P3-injected brains. Peak density of retrogradely labelled cells in claustrum was reached during the second postnatal week. (j) Mean±sem of all retrogradely labelled claustrum cells counted in every 5^th^ section along the AP extent of the claustrum of a given brain. Individual brains shown as circles. There was a steady increase in the number of back-labelled claustrum cells with increasing age when using carbocyanine dye tracing, but this did not reach statistical significance (Spearman’s r=0.9, p=0.0833). (k) Mean±sem of all retrogradely labelled claustrum cells counted in every 10^th^ section along the AP extent of the claustrum of a given brain. Individual brains shown as triangles (2-day survival) and squares (5-day survival). There was no significant difference between two- and five-day survival following CTB injections (ANOVA, F(6,15)=1.606, p=0.2134). There was a strong positive correlation between the number of back-labelled claustrum cells and increasing brain age (Spearman’s ρ=0.7857, p=0.048). (l) Anterior claustrum projects more strongly to ACA throughout development. Retrogradely labelled cell counts were binned by anterior-posterior location. ‘Very anterior’ corresponds to slices at and just posterior to the beginning of striatum. ‘Anterior’ corresponds to slices at and just posterior to the first midline crossing of the corpus callosum. ‘Middle’ is at the level of the anterior commissure midline crossing, and ‘posterior’ is caudal to that, up to and including the caudal end of claustrum. More ACA projecting neurons are located in the anterior claustrum than in the more posterior regions. There was a significant effect of both age at injection (ANOVA, F (6, 66)=4.537, p=0.0007) and anterior-posterior position (F (3, 66)=17.53, p<0.0001). Scale bars = 100µm and 1mm (insets).

For each brain, a carbocyanine crystal was placed in the approximate location of ACA from the dorsal surface or the midline of intact hemispheres, and brains or hemispheres were incubated for 4-6 weeks to allow the dye to diffuse. Location of the injection site was determined to be mostly in secondary motor cortex (MOs), with some crystal placement sites extending into ACA, prelimbic area (PLA) or laterally into primary motor cortex (MOp). With the exception of the youngest brains, we always found some back-labelled cells in the claustrum or dorsally adjacent cortex (Fig 5). Following ACA/MO dye placements, we observed a fibre plexus in lateral cortex that may have included the dorsal claustrum or might be just dorsal to it. Cells in this fibre plexus were included in the cell counts, so the carbocyanine data presented here is possibly an overestimate of the early connectivity between claustrum and anterior cingulate cortex. ACA/MO-projecting neurons in the putative claustrum were counted in every 5^th^ coronal section along the anterior-posterior extent of the claustrum, starting at the level of the anterior-most striatum. There was an increase in the number of ACA-projecting, putative claustrum cells with increasing brain age, but this did not reach statistical significance (Spearman’s r=0.9, p=0.0833). At all ages investigated using carbocyanine dye tracing, the majority of back-labelled neurons were found in the more anterior slices.

Claustrum neurons projecting to ACA have been reported to co-project to orbitofrontal cortex (OFC) but not primary motor or sensory cortices in adult mice (Chia et al. 2020). Thus, crystal placement sites extending into M1/M2 are likely to label a different population of cells to those selective to ACA.

To extend the time-line, ensure inclusion of only claustral neurons, and improve targeting of injection site to ACA, we switched to an *in vivo* CTB-injection strategy. CTB was injected into ACA of CD1 pups and mice based on scaled stereotaxic coordinates (>P7, n=32 mice) or head surface landmarks (<P5, n=23 mice), and the injection site was confirmed to include ACA for each brain included in the analysis here (n=22). To ensure that we only included cells located in the unambiguously identifiable claustrum, we restricted our cell counts to sections at the level of the striatum. The youngest age with on-target CTB-injections into ACA was P3, and we could identify a small number of retrogradely-labelled claustrum neurons in both brains (n=2; Fig 5F,K). Many more back-labelled neurons were found in the claustrum of mice injected at P8 and P10 (n=5 and 6 respectively). There was no further increase in the number of back-labelled cells after P10, suggesting that adult-like innervation density is reached within a week of the first claustral axons reaching ACA. ACA-projecting neurons in the putative claustrum were counted in every 10^th^ coronal section along the anterior-posterior extent of the claustrum, starting at the level of the anterior-most striatum. As there was no significant difference between two and five-day post-injection survival (ANOVA, F(6,15)=1.606, p=0.2134; Fig 5K), we grouped the data only by injection age for all subsequent analysis. There was a strong and significant positive correlation between brain age and number of ACA-projecting claustrum neurons with increasing age (Spearman’s r=0.7857, p=0.048).

There was a pronounced anterior-posterior gradient, with many more back-labelled cells found in the anterior claustrum at most ages. To analyse this further, we binned cell counts into four anterior-posterior positions (Fig 5L). There was a significant effect of both age of brain at injection (ANOVA, F (6, 66)=4.537, p=0.0007) and anterior-posterior position (F (3, 66)=17.53, p<0.0001). From P14 onwards, the highest number of cells was always located in the ‘very anterior’ claustrum, which corresponds to sections at the level of the anterior end of the striatum. Following P8 and P10 injections, the highest number of cells was found in the ‘anterior’ claustrum bin, which corresponds to sections at approximately the level of the anterior-most corpus callosum midline crossing. In the youngest brains there were so few cells, that no gradient was evident. Overall, there was no significant difference between ‘very anterior’ and ‘anterior’ claustrum, but both were significantly different from ‘middle’ and ‘posterior’ claustrum (Tukey’s multiple comparison test: very anterior vs anterior p=0.9821, very anterior vs middle p=0.0005, very anterior vs posterior p<0.0001, anterior vs middle p=0.0003, anterior vs posterior p<0.0001, middle vs posterior p=0.2816).

## Discussion

The claustrum is notoriously difficult to delineate in brains without a prominent extreme capsule, and this problem is compounded in developing brains. Commonly used strategies to delineate the adult claustrum, such as the parvalbumin-plexus, retrograde labelling from retrosplenial cortex, and regional differences in myelination are all suitable for use after the third postnatal week in mice. Moreover, they can be useful in combination to distinguish subregions within the adult claustrum (Grimstvedt et al. n.d.). We have demonstrated here that none of them can be used to demarcate the claustrum during the first two postnatal weeks. Nurr1 has previously been used at the transcriptomics level as a marker for adult claustrum cells (Niu et al. 2022), and when comparing claustrum in different species (Montiel et al. 2011; Wang et al. 2011; Puelles 2014; Binks et al. 2019), however, it is widely expressed in lateral cortex (Arimatsu et al. 2003; Hoerder-Suabedissen et al. 2009), and thus not one of the preferred claustrum labelling strategies in adult brains. There is, however, a particularly dense patch of Nurr1-immunoreactive cells evident in the claustrum. We have shown that this patch of high Nurr1+ cell density overlaps with the cluster of RSP-projecting claustrum cells, and is mostly contained within the ‘bird’s nest’-like arrangement of myelinated fibres surrounding the ventral claustrum. The Nurr1+ claustrum patch is distinct, even in neonatal mouse brains. It is present along the anterior-posterior extent of the claustrum, with subtle variations in shape, size and relative cell density, and contrasts with the much lower Nurr1 expression levels observed along the cortical subplate. Nurr1 immunohistochemistry with the protocol used here is suitable for different species including pigs, ferrets and human when labelling subplate neurons (unpublished observation; Montiel et al. 2011; Wang et al. 2011; Molnár and Clowry 2012), and may thus provide a good starting point for comparing claustrum development across different species.

Here we report that the claustrum has a short period of peak neurogenesis centred on E12.5 in the mouse. This falls within the time window of claustrum neurogenesis that can be inferred from the diagrams in Smart and Smart (1977), but narrows the time frame (Smart and Smart 1977). This period of peak neurogenesis is the same as the amygdala (Clancy et al. 2001), which shares a similar anatomical position to the claustrum, as well as other developmental similarities (Butler and Molnár 2002; Molnár and Butler 2002). In contrast, the dorsal and ventral neighbours of the claustrum, subplate and endopiriform nucleus respectively, both have earlier peak neurogenesis (data shown here and Hoerder-Suabedissen and Molnár 2013).

The neurons of the ventrally adjacent endopiriform nucleus are generated earlier (at E11.5). BrdU labelling alone enabled a rough distinction between claustrum and endopiriform nucleus, based on their distinct birth dates at E12.5 and E11.5, respectively, which may be useful in future studies, and highlighted that the birthdates of subplate and endopiriform nucleus are more similar to each other than to the claustrum. We did not include a separate region of quantification for the dorsal-most part of endopiriform nucleus (often referred to as ‘medial endopiriform nucleus’ in adult mouse brains), as there were not enough landmarks to clearly delineate its boundaries in developing brains. However, during quantification it was observed that the region of dense E12.5-generated BrdU+ cells extends a short distance ventral to the MBP-sparse claustrum region, whereas the region with the most E11.5-generated BrdU+ cells extends slightly dorsal to the region used for endopiriform nucleus quantification. Thus, there appears to be a ventro-dorsal gradient of neurogenesis within the endopiriform nucleus, with more ventral cells being generated earlier in development. This result is similar to that reported for neonatal rats (Fang et al. 2021).

Is there significant cell death in claustrum during the postnatal period? The density of Nurr1+ cells in claustrum is much higher in P5 and P8 brains compared to older brains, but the cross-sectional area of the claustrum as determined from the outline of the Nurr1+ patch does not increase significantly with age. This may suggest a high degree of cell death occurring in claustrum towards the end of the first postnatal week, but further investigation, such as normalisation to a pan-neuronal marker, would be required to confirm this. Moreover, the brain also expands in the anterior-posterior dimension, which may change the volume of the claustrum, resulting in reduced cell density in the absence of cell death.

We provide evidence for the presence of a subplate/layer 6b medial to the claustrum. Layer 6 markers surround the claustrum. *Crym*, present dorsally in both layer 6a and 6b, surrounds the claustrum on all sides, but is absent from the claustrum itself (Brigman et al. 2009; Wang et al. 2017). In other work, we found that *Rprm* labels layer 6a throughout lateral cortex and is found exclusively lateral to the claustrum, in layer 6 of insula (Grimstvedt et al. n.d.). *Ctgf*, a subplate/layer 6b marker (Heuer et al. 2003; Hoerder-Suabedissen et al. 2009), is absent from the claustrum itself, but present both in endopiriform nucleus, as well as medial to the claustrum (Grimstvedt et al. n.d.; Erwin et al. 2021). *Nr4a2*/Nurr1, on the other hand, is present at low levels in the subplate/layer 6b (Hoerder-Suabedissen et al. 2009; Hoerder-Suabedissen and Molnár 2013) but expressed very strongly and densely in the claustrum (this work; Arimatsu et al. 2003; Fang et al. 2021). Cplx3+ cells, *Ctgf* expression and E11.5-born neurons, all typical of subplate neurons, are present medial to the claustrum in our and other’s work. Despite the co-expression of Nurr1 in claustrum and subplate, its varying levels of expression, the contrasting expression patterns of *Ctgf* and *Cplx3* and other genes, the distinct birth-dates and significant differences in the time-course of formation of connections all distinguish these structures. In summary, we suggest that there is a subplate/layer6b medial to the claustrum.

One of the most striking features of the adult claustrum is its dense, and often reciprocal, connectivity with the rest of the cerebral cortex. Understanding this connectivity architecture represents a crucial foundation upon which we can then develop a deeper understanding of the role of the claustrum in brain function. The claustrum contains many distinct groups of neurons in terms of outgoing projections (Zingg et al. 2018; Chia et al. 2020; Marriott et al. 2021; Peng et al. 2021; Chevée et al. 2022), and may thus be recruited for different functions. Although the mature claustrum may be recruited during a variety of brain states or behaviours in adults, behaviours or brain states displayed before claustro-cortical connectivity emerges are improbable primary functions of the claustrum. Thus, we specifically investigated when claustrum axons first connect to two distinct ipsilateral target regions: anterior cingulate and retrosplenial cortex. RSP projections were chosen, because they are widely used in adult brains as a means to access claustrum cells for genetic manipulation via retrograde viral transfection including in species other than mouse (Zingg et al. 2018; Dillingham et al. 2019; Marriott et al. 2021; Shelton et al. 2022), and ACA because of its reported dense innervation by claustrum (Wang et al. 2017; White et al. 2018; Zingg et al. 2018; Chia et al. 2020).

The profile of RSP projecting cells through development we report here is consistent with the tight cluster of cells observed in the claustrum in the adult (Zingg et al. 2018; Marriott et al. 2021; Shelton et al. 2022). We find no projections to RSP from insula and other claustrum adjacent structures even in the youngest brains. Once claustrum neurons project to RSP in large numbers (e.g. from approximately P16), retrograde tracing from RSP becomes a suitable claustrum delineation and genetic access strategy also in developing brains. We find claustrum cells retrogradely labelled from ACA, before we find cells labelled from RSP. Retrograde tracing from ACA, however, cannot be used in early development for selective access to the claustrum, because we found nearby non-claustrum cells retrogradely labelled from ACA at all ages investigated.

Claustrocortical axons develop late relative to other key cortical pathways, and in particular the claustral projections to RSP emerge late relative to those targeting ACA. There is no further increase in the number of cells projecting to ACA after P10, suggesting that axon outgrowth and targeting to this region is complete by P10. Conversely, the number of claustrum cells with axons to RSP continues to increase until at least P16, suggesting that this is an even more slowly maturing pathway. In contrast, local, subplate, interhemispheric, corticothalamic and thalamocortical connections all form earlier in development. Local cortico-cortical connections in the mouse have been reported from E16.5 onwards (Huffman et al. 2004). We observed retrogradely labelled subplate cells for all of our carbocyanine placements, including all P2 brains, as has been reported previously (Hoerder-Suabedissen and Molnár 2012). Similarly, interhemispheric projections in the rat have been demonstrated to be present shortly after birth for callosally projecting neurons (Ivy and Killackey 1981; Olavarria and van Sluyters 1985). Cortico-thalamic L5 axon collaterals in mice are present in their thalamic target regions by P2, although axonal arbours continue to elaborate for several more days (Grant et al. 2016; Hoerder-Suabedissen et al. 2018). Thalamocortical axons first reach the cortex during embryonic development (Molnár and Blakemore 1995; Higashi et al. 2002; Antón-Bolaños et al. 2019), and have segregated into their periphery-related pattern in layer 4 by P4 (Agmon et al. 1995). This comparison with the maturation of other cortical axon pathways highlights that claustro-cortical projections start to develop, and are most elaborated after similar processes in other major cortical pathways have been underway for some time. This may suggest that the claustrum is primarily involved in ‘higher cognitive functions’ that become necessary only once animals have to fend for themselves.

In adult brains, a considerable proportion of claustrum neurons projects to both ACA and RSP (Zingg et al. 2018), but our data does not permit us to identify whether a single neuron first extends an axon into ACA, before innervating RSP. This could be addressed in the future by dual-colour tracing from both regions.

Cortical areas often project bilaterally to claustrum, but reciprocal connections from claustrum back to cortex are almost exclusively ipsilateral (Smith and Alloway 2010, 2014; Smith et al. 2012; Zingg et al. 2014; Wang et al. 2017; Shelton et al. 2022). Thus, at most ages we only analysed ipsilateral projections during development, to facilitate finding any cells at all, at ages when only few claustrum neurons are retrogradely labelled.

In isolation, projections to two cortical target regions do not permit us to rule out any one claustrum function. It is intriguing, however, that delta waves, the electroencephalogram (EEG)/electrocorticogram (ECoG) hallmark feature of slow-wave sleep, are undetectable in young rats until P11 (Jouvet-Mounier et al. 1969; Gramsbergen 1976; Frank and Heller 1997; Seelke and Blumberg 2008). By P18, the EEG during different behavioural states is indistinguishable from that of adult rats (Gramsbergen 1976). Similarly for mice, slow waves first emerge between P9 and P12 (depending upon mouse strain), and by P12 the EEG during slow wave sleep is indistinguishable from that of adult mice (Daszuta and Gambarelli 1985). Thus, it is reasonable to assume that mouse cortices would also start to exhibit delta wave EEG activity from around P11, or possibly a few days earlier. We observed that claustrum to ACA projections reach their full number at around this time-point. Although we did not test for synaptic connection strength, the time-frames of delta-wave EEG emergence and claustro-cortical connectivity to ACA provide some evidence that the establishment of claustrum connectivity is necessary for the developmental emergence of cortical slow-wave activity. Additionally, the time-frame of claustro-cortical connectivity emergence overlaps with other major changes in mouse pup behaviour, such as the emergence of active sensory processing (i.e. active whisking, and eye opening).

In summary, it is a challenge to probe claustrum using adult markers as many of these are not expressed early, however we found a strategy to reliably identify the claustrum throughout the postnatal period. Our combined lines of investigation allow us to draw firm conclusions about the temporal origin of the claustrum and its innervation of overlying cortex. Claustrum neurons in the mouse are generated across a short time-window centred on E12.5, which is later than the peak neurogenesis in the neighbouring subplate and endopiriform nucleus. Innervation of two of the claustrum’s main adult cortical target regions – anterior cingulate and retrosplenial cortex – emerge with distinct developmental trajectories. Projections to ACA emerge earlier and progress to an adult-like innervation pattern faster than projections to RSP, and this may have a bearing on the emergence of EEG-measurable slow-wave sleep patterns.

## Acknowledgements

This project was funded by the Wellcome Trust (A.M.P.), the European Research Council (ERC) under the European Union’s Horizon 2020 research and innovation programme (grant agreement No 852765; A.M.P.) and the Medical (G0900901; Z.M. & A.H-S.) and Biotechnology and Biological Sciences (BB/P003796/1; S.B.) Research Councils. A.M.S was funded by a Clarendon Fund graduate scholarship, G.O.S. was in receipt of a fellowship from the “la Caixa” Foundation (ID 100010434; fellowship code LCF/BQ/EU20/11810053), T.H.D. received generous support from Lincoln College (Oxford) and Z.M is supported by the “Einstein Stiftung” (Berlin).

## Supplementary Material

**Supplementary Figure 1:**
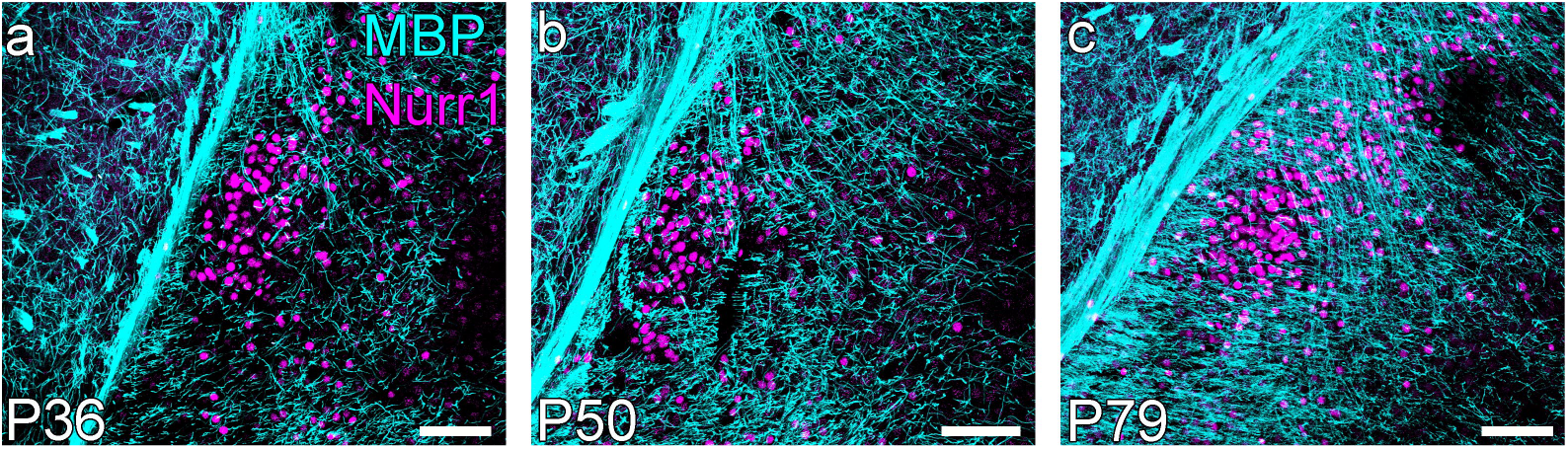
At the level of the anterior commissure, the Nurr1+ patch of claustrum cells sits in the centre of the MBP+ ‘bird’s nest’ of myelinated fibres surrounding the claustrum. (a-c) Maximum intensity projection confocal laser scanning microscope images of the claustrum region at different postnatal ages, immunohistochemically stained for Nurr1 and MBP. Scale bars = 100µm

**Supplementary Figure 2:**
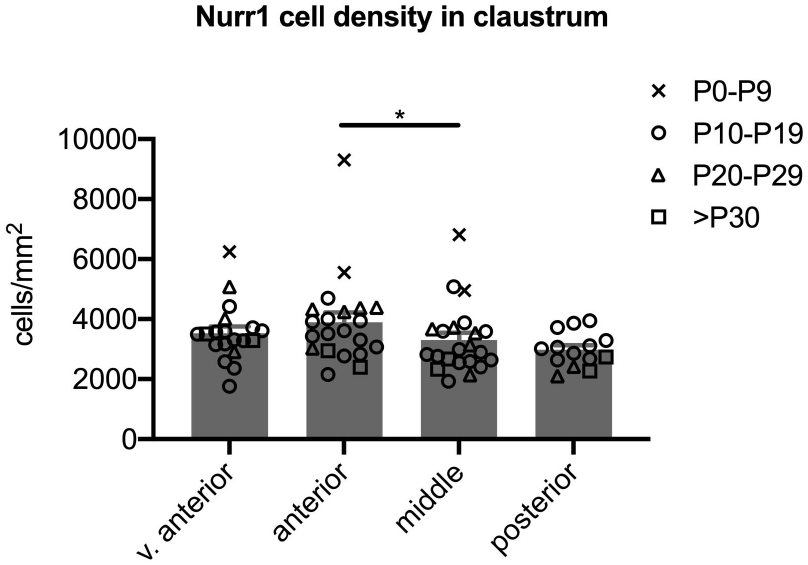
Density of Nurr1+ cells in the claustrum is much higher during the first ten days after birth, and stable thereafter. Nurr1+ cell density was quantified in three or four sections for each brain (n=22 brains aged P5-P36). There is a strong effect of age on Nurr1+ cell density in the claustrum (main effects ANOVA F (9, 63) = 16.55, p<0.0001), which is exclusively due to the much higher cell density throughout the claustrum in the youngest (and therefore smallest) two brains (Tukey’s multiple comparison test p<0.01 for all comparisons involving P5 and P8 brains; data represented by ‘x’ in the graph). There is also a significant effect of anterior-posterior position on Nurr1+ cell density within the claustrum (main effects model ANOVA F (3, 63) = 3.701, p=0.0161). Tukey’s multiple comparison test indicates that only the Nurr1+ patch density at the level of the anterior commissure midline-crossing (‘middle claustrum’) is significantly different from the anteriorly adjacent claustrum (Tukey’s multiple comparison test, p=0.0318). * p<0.05

